# An isothermal method for sensitive detection of *Mycobacterium tuberculosis complexes* using CRISPR/Cas12a *cis-* and *trans-*cleavage

**DOI:** 10.1101/2020.02.03.933101

**Authors:** Haipo Xu, Xiaolong Zhang, Zhixiong Cai, Xiuqing Dong, Geng Chen, Zhenli Li, Liman Qiu, Lei He, Xiaolong Liu, Jingfeng Liu

## Abstract

Tuberculosis is still one of the most serious infectious diseases resulting in lethal death worldwide. The traditional method is still not enough to meet the clinical requirements of rapid diagnosis, high specificity and sensitivity. Fast, sensitive and accurate detection of *mycobacterium tuberculosis* (MTB) is an urgent need for the treatment and control of tuberculosis disease. Clustered Regularly Interspaced Short Palindromic Repeats (CRISPR)-associated proteins (Cas12a) exhibits strongly nonspecific degradation ability of exogenous single-strand nucleic acid (trans-cleavage) after specific recognition of target sequence. We purified Cas12a protein and selected a proper guide RNA (gRNA) based on conserved sequences of MTB from gRNA library we designed. Then, we proposed a novel method based on recombinase polymerase amplification (RPA) and CRISPR/Cas12a nuclease system for specific and sensitive detection of MTB DNA. The assay based on fluorescence detection pattern showed 4.48 fM of limit of detection (LOD) and good linear correlation of concentration and fluorescence value (R^2^=0.9775). Also, it showed good performance in distinguishing other bacteria. Furthermore, its clinical performance was evaluated by 193 samples and showed sensitivity of 99.29% (139/140) and specificity of 100% (53/53) at 99% confidence interval, respectively, compared with culture method. The CRISPR/Cas12a system showed good specificity, excellent sensitivity and accuracy for MTB detection, and it meets requirements of MTB detection in clinical samples and has great potential for clinical translation.

---

Tuberculosis (TB), which is caused by *Mycobacterium tuberculosis* (MTB) and could spread through aerosol in the air, for example by coughing, remains one of the top 10 causes of death and the leading cause of death from a single infectious agent (ranking above HIV/AIDS) worldwide ^1^. Therefore, the fast, sensitive and accurate diagnosis of active TB and drug resistance (for example, rifampicin, isoniazid, pyrazinamide) is crucial both for the individual and population level to reduce morbidity, mortality and transmission among patients. Sputum smear microscopy is the primary test for TB diagnosis, but it is significantly limited by its laborious detection process and relatively low detection sensitivity of only 56% to 68%^2, 3^. In addition, the microscopy performs poorly on HIV co-infected specimens because the amount of MTB is low in sputum specimen. Meanwhile, it cannot distinguish MTB and *non-tuberculosis mycobacteria* (NTM) which of patients are treated totally differently due to different pathogenicity and drug susceptibility profiles ^4–6^, although they show similar clinical manifestation. So far, culture-based mycobacterial detection method is still the gold standard for the laboratory diagnosis of tuberculosis due to its high sensitivity, specificity and accuracy^7^. But, it needs extensive laboratory experiences and usually takes up to 4-6 weeks to achieve satisfied results, which does not meet the rapid test requirement, and it significantly hindered the prompt treatment and isolation patients which is crucial for disease control. Moreover, it exists the cases that some bacilli cannot be successfully cultured, which might arise error results ^8^.

Genetic sequencing of MTB DNA was reported to be a more accurate strategy for TB diagnosis. The Xpert MTB/RIF system (Cepheid, USA), a polymerase chain reaction (PCR) based test, is the only one accepted by WHO from 2010 for detecting TB infection and rifampicin resistance. Though it has many advantages including full automation, integration and better accuracy than microscopy or culture methods, its sensitivity for some bacilli clinical specimen is still inadequate. Furthermore, the cost of cartridges, instrumentation and maintenance of Xpert MTB/RIF system make it impossible to set-up in many high TB burden countries, which further limited its usefulness^9, 10^.

In recent years, isothermal amplification assays using loop mediated amplification (LAMP) ^11^, helicase dependent amplification (HDA) ^12^ and recombinase polymerase amplification (RPA) ^13–15^ have been extensively applied in the field of molecular diagnosis including pulmonary TB diagnosis ^16–18^. Especially, the RPA reaction which replaces the thermal cycles needed for PCR reaction and works at temperatures between 25°C and 42°C with three core enzymes, permits rapid DNA amplification in minutes. Recently, the RPA based methods have been used for detection of MTB and other pathogen, while it always had high false positive results in clinical samples^19^. To further improve specificity, long oligonucleotide probe with fluorophores and quenchers or other modifications of probe, such as biotin, tetrahydrofuran, etc, have been designed to recognize RPA amplicon for specific cleavage by nucleases, and these newly developed methods have been successfully applied for pathogen diagnosis^20–22^, liquid biopsy^23^, and SNP/genotyping^24^. However, the synthesis of long oligonucleotide probes with multiple modifications is cost, and different probes with the same amplified region may also affect the efficiency of RPA. Especially, the base mismatches near the middle of probes or the 3’-OH end of primers might significantly affect the efficiency of RPA reaction, which could lead to reduced amplification efficiency and false positive results ^19, 25^.

Microbial Clustered Regularly Interspaced Short Palindromic Repeats (CRISPR)-associated (CRISPR-Cas) adaptive immune system, consists of guide RNA(s) for target recognition and a Cas enzyme for target cleavage^26–29^. The sequence-specific recognition capabilities of the CRISPR-Cas system make it as a promising genome engineering tool for genome editing^29, 30^, cancer target therapy ^31^, and gene therapy^32^. Recently, the CRISPR-Cas nuclease systems (Cas12, Cas13 and Cas14)^26, 33–35^ also have been developed for nucleic acid detection with high sensitivity and specificity. These CRISPR-Cas nuclease systems use the gRNA to recognize the RNA (Cas13a and Cas13b) or DNA (Cas12a and Cas14) targets, then followed by activating cleavage capacity of Cas nuclease to degrade multiple foreign nucleic acid reporters for target detection. Combining with nucleic acid amplification strategy, the Cas nucleases systems have been used for highly sensitive and specific detection of nucleic acids and became as powerful tools for developing molecular diagnostic techniques^26, 33, 34^.

To achieve prompt, sensitive and specific confirmed diagnosis of MTB, herein we combined the RPA reaction with the CRISPR-Cas nuclease (Cas12a) system to develop a novel MTB DNA detection method (Fig. 1). Firstly, the sequence of MTB with protospacer adjacent motif structure (5’-TTTN-3’) for Cas12/guide RNA (gRNA) recognition was selected for gRNA design. The gRNA consists of two parts, universal spacer sequence (5’-UAAUUUCUACUAAGUGUAGAU-3’) at the 5’ end (also near the PAM structure end) and specific sequence at the 3’ end. And gRNAs were selected via evaluating by multiple softwares, such as CRISPR-DT, NUPACK, Nucleotide BLAST etc. Then the feasibility of Cas12a/gRNA trans-cleavage reaction was validated by the purified Cas12a. Subsequently, the suitable amplicon screening was performed and tested to effectively avoid false amplification of RPA. In the Cas12a/gRNA trans-cleavage fluorescence for MTB assay, in the presence of MTB DNA, they could be amplified by RPA reaction. Later, through Cas12a/gRNA complexes binding to amplified MTB DNA, Cas12a mediated nonspecific degradation of nonspecific fluorescent probes, which were polythymidine oligonucleotides with fluorophore at the 5’ end and quencher at the 3’ end as MTB DNA reporters, could release the fluorophores from the quenchers to increase fluorescence intensities for detection. Here proposed a novel isothermal method based on RPA and CRISPR-Cas system which has very low limitation on MTB detection in our model testing system, such as 4.48 fM of LOD, range over 6 orders of magnitude (R^2^=0.9775), and could be applied to practical applications on clinical samples with excellent specificity, sensitivity and accurate, which are comparable with the result to the gold standard of culture method.

**Figure 1.**
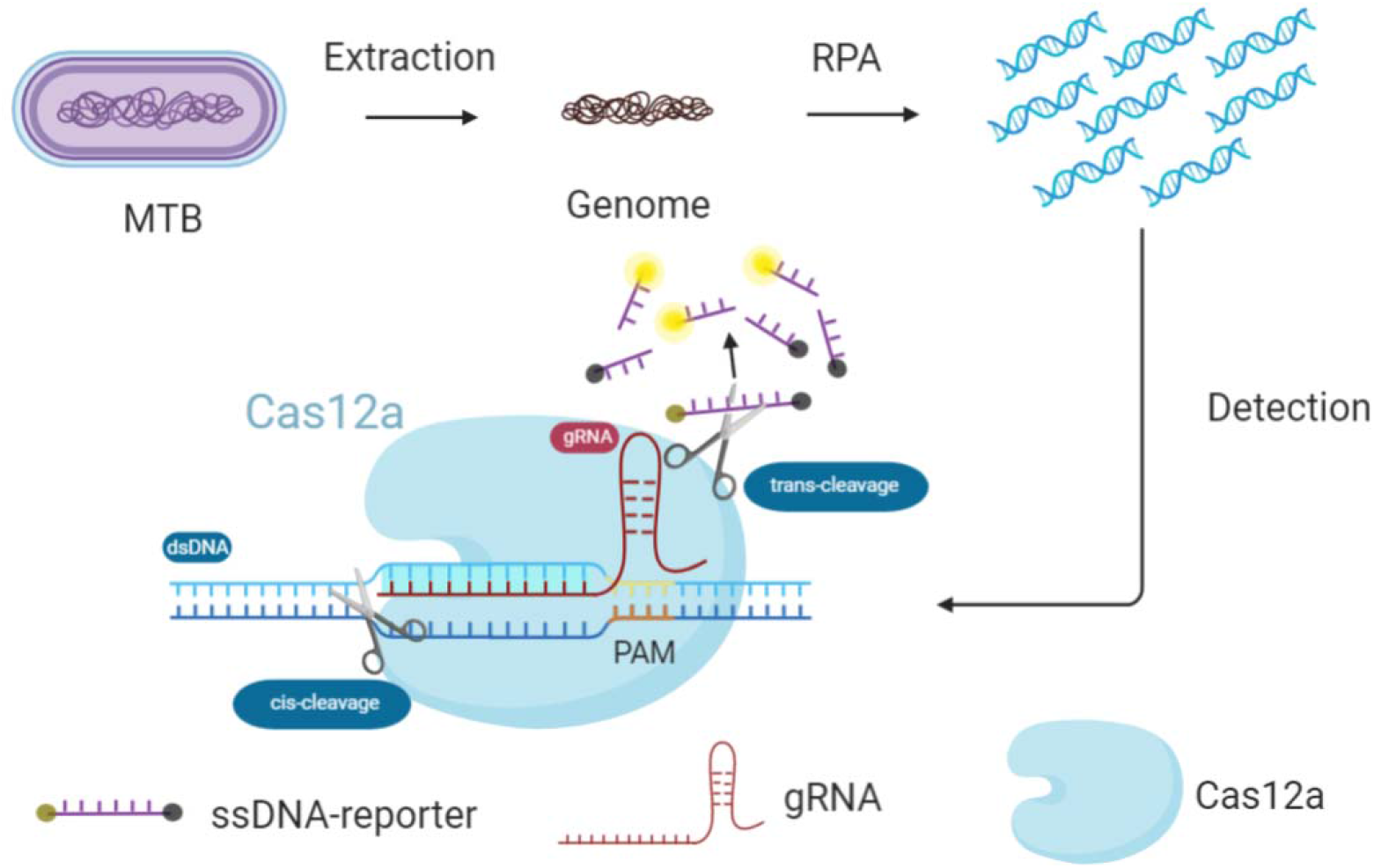
Schematic diagram of the Cas12a/gRNA trans cleavage fluorescence signal amplification system for MTB detection. Genome of mycobacterium tuberculosis complex (MTB) was extracted into RPA reaction for amplification. Positive fluorescent signals were captured when probes were cleaved by activated Cas12a under target with PAM sequence at 5’ end that recognized by gRNA.

## Materials and Methods

### Target Selection and gRNA Design for MTB-Cas12a System

MTB-specific fragment of the insertion sequence IS1081, which exists in all *mycobacterium tuberculosis complexes*^36^, was selected as the target sequence for design of gRNA to target MTB and RPA reaction primers. To obtain the suitable guide RNA for MTB, gRNAs targeting IS1081 dsDNA were designed and scored by CRISPR-DT online software for Cpf1 (Cas12a) (http://bioinfolab.miamioh.edu/CRISPR-DT/interface/Cpf1_main.php), and target efficiency less than 0.6 was excluded. Meanwhile, the second structure and homology of remaining gRNAs were further evaluated by NUPACK (http://www.nupack.org/) and Nucleotide BLAST (https://blast.ncbi.nlm.nih.gov/Blast.cgi?PROGRAM=blastn&PAGE_TYPE=BlastSearch&LINK_LOC=blasthome), respectively. Afterwards, RPA primers were designed through Primer Premier 5.0 software and selected using agarose gel electrophoresis according to the design manual of TwistAmp Assay (TwistDx Ltd., England) https://www.twistdx.co.uk/docs/default-source/RPA-assay-design/twistamp-assay-design-manual-v2-5.pdf?sfvrsn=29.

All RNA, IS1081 or IS6110 and modified DNA oligo were synthesized by General Biosystems (Anhui, China). Plasmids were obtained from the construct that the insert sequence IS1081 (Genbank ID: CP003248.2) and IS6110 (Genbank ID: X17348.1) were put into vector PUC57 respectively. Sequence information can be seen in the Supplement Data. The fluorescent polyoligonucleotides probe (ssDNA-FQ) was labeled with a fluorescent dye at the 5’ end and a quenching group at the 3’ end then purified with HPLC. The fluorescence could be detected only when the fluorescent dye and quenching group were separated. Forward and Reverse primers for RPA and non-specific single strand DNA (ssDNA, non-target) for Cas12a/gRNA trans cleavage reaction were obtained from Sangon Biotech (Shanghai, China) with PAGE purification. All DNA/RNA oligonucleotides and plasmids were diluted with RNase-free water and the concentrations were determined by Nanodrop 200 (Thermo Fisher, USA). All of these were stored at −20°C until use. The sequences of DNA/RNA oligonucleotides used in the assay were shown in Table 1.

**Table 1.**
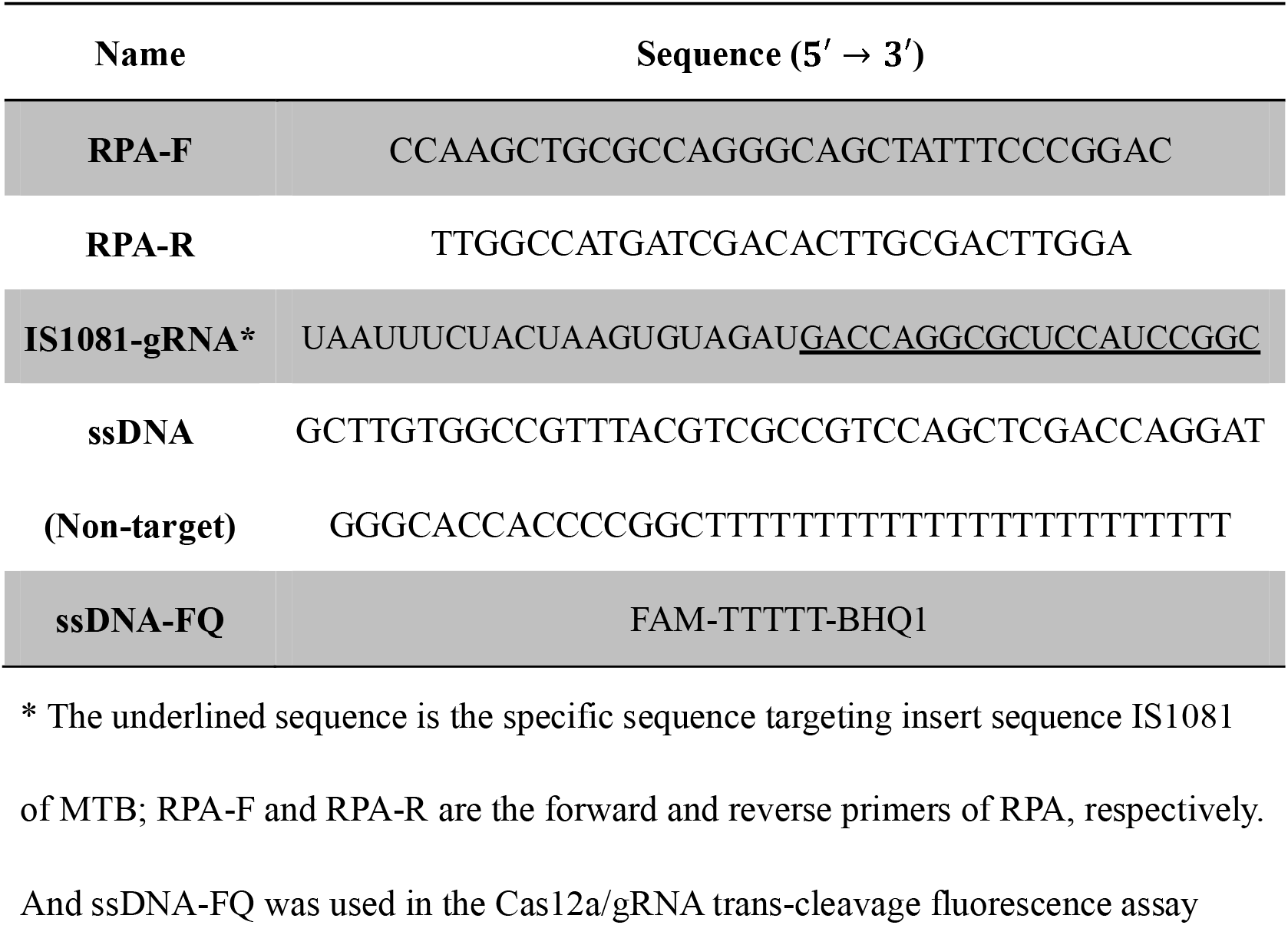
The oligonucleotide sequences of RPA primers, gRNA and non-specific single strand DNA for MTB Cas12a/gRNA trans-cleavage assay

### Feasibility of Purified Cas12a for MTB Detection

The DNA plasmid encoding Cas12a (also named Cpf1) with 6×His-tag, maltose-binding protein (MBP) and TEV protease cleavage site was selected for LbCas12a expression. The plasmid 6His-MBP-TEV-huLbCpf1 (plasmid # 90096) (see Figure.1 and Supplementary M3 in the Supplement Data) was purchased from Addgene. After extracted from amplified strains, the plasmids were transformed into BL21(DE3) pLysS Chemically Competent Cells (TransGen Biotech). Then transformed strains were recovered from Luria-Bertani solid growth media and turned to 5 mL Luria-Bertani liquid growth media. After that, the LbaCas12a protein was expressed and purified according to Chen’s method ^26^. Briefly, a 5 mL starter culture was grown in Luria-Bertani liquid growth media (LB, containing of10 g/L tryptone, 5 g/L yeast extract, 10 g/L NaCl, pH 7.0, Sigma) at 25°C overnight, and then further transferred into 1 L of LB and inoculated for growth at 37°C under 300 rpm shaking until the OD600 approximately reach to 0.6. Afterwards, all bacterium was cooled down and supplemented with 500 μM isopropyl-1-thio-b-D-galactopyranoside (IPTG, TransGen Biotech) at 16°C for 16 h to allow protein expression. After centrifugation at 5200 rpm for 30 min at 4°C, the bacterial precipitation was harvested and stored at −80°C until use.

The protein purification was performed at 4°C. Firstly, the bacterial precipitation was crushed and lysed by sonication in Lysis buffer (50 mM Tris-HCl, 500 mM NaCl, 5% (v/v) glycerol, 1 mM TCEP, 0.5 mM PMSF and 0.25 mg/mL lysozyme, pH 7.5). After centrifugation, the supernatant was filtered via a Stericup 0.22 μm filter (EMD Millipore). The filtered supernatant was pipetted to Ni-NTA resin (GE) for 30 min, and washed with lysis buffer. Then, the resin was resuspended in 1×TEV digestion buffer (50 mM Tris-HCl, 50 mM NaCl, 0.5 mM EDTA, 1 mM DTT, pH 8.0) with 250 units of TEV protease (Beyotime Biotechnology) overnight at 4°C with rotation. To obtain purer proteins, the steps on Ni-NTA resin was repeated. The protease treated LbCas12a was dialyzed in Buffer A (20 mM Tris-HCl, 125 mM KCl, 5% glycerol, 1 mM TCEP, pH 7.5) for three times at 4°C, and further purified on a 5 mL HiTrap SP HP sepharose column (GE Healthcare). After washing with three column volumes of Buffer A, LbCas12a was eluted using a linear gradient from 0-100% Buffer B (20 mM Tris-HCl, 1 M KCl, 5% glycerol, 1 mM TCEP, pH 7.5) over 20 column volumes. The eluted proteins were further purified on Superdex 200 column (GE Healthcare) with elution Buffer (20 mM Tris-HCl, 200 mM KCl, 5% glycerol, 1 mM TCEP, pH 7.5). The existence of LbCas12a protein was confirmed by SDS-PAGE and Coomassie Blue staining, and then mixed with equal volume of glycerol after quantification and stored at −20°C for further usage. To test the *cis-* and *trans-*cleavage activity of purified LbCas12a protein and the feasibility of Cas12/gRNA trans cleavage for MTB detection, we were observing the alteration of amount and integrity of substrates and nonspecific single strand DNA by 4% agarose gel electrophoresis and the fluorescence intensity was collected by detector with/out target plasmid, gRNA or non-target ssDNA. The Cas12a/gRNA trans-cleavage for MTB detection assay was as follows: 50 nM LbCas12a incubating with 36 nM IS1081-gRNA, 80 ng IS1081 plasmid and 50 nM non-target ssDNA (replaced with ssDNA-FQ in Cas12a/gRNA fluorescence assay) in 1× reaction buffer (50 mM NaCl, 10 mM Tris-HCl, 10 mM MgCl_2_, 100 μg/mL BSA, pH 7.9), incubated at 37°C for 2 h.

### Mycobacterium tuberculosis (MTB) Detection by Cas12a Fluorescence Assay

To improve signal amplification efficiency and simplify operation, 1 μL of DNA extracted from amplified clone strains as template was added into RPA reaction mixture of 12.5 μL volume. RPA reaction mixture including 0.24μM~0.72μM forward and reverse primers, 1.8 mM dNTP, 1×Reaction Buffer, 1×Basic E-mix, 0.625 μL 20×Core Reaction Mix, 1 μL genome, and 14 mM MgOAc and DEPC water up to 12.5 μL and incubation at 37°C~42°C for 1 h. According to the design manual of TwistAmp (TwistDx Ltd., England) assay, multiplex sets of RPA primers were selected as candidates, including primers from published literature^18, 37^ (see Table.1 in the Supplemental Data) and tried to get best conditions from RPA reaction *via* optimization for concentration of dNTP, Mg^2+^, primers and reaction temperature with TwistAmp® Liquid Basic/Basic RT (TwistDx, England).

After RPA reaction finished at one hour, 7.5 μL Cas12a/gRNA mixtures of 36 nM gRNA, 50 nM ssDNA-FQ, 50 nM Cas12a in 1× reaction buffer was added into the cap of RPA reaction tube to the final volume of 20 μL. Then transient high-speed centrifugation was performed to mix RPA productions with Cas12a/gRNA system. The Cas12a/gRNA trans-cleavage fluorescence reaction was incubated at 37°C for 2 hours and its fluorescence was collected every 1.25 minutes by fluorescence detector (Applied Biosystems 7500 Real-Time PCR System in this study) during reaction.

As we defined, positive result of sample was refined as that the fluorescence intensity collected by detector was 1.5 times more than that of negative control. Others were negative values of samples. Negative control was non-MTB DNA as template for Cas12a/gRNA trans-cleavage fluorescence assay.

### Performance of MTB-Cas12a Fluorescence Assay

For evaluation the sensitivity of Cas12a/gRNA for MTB fluorescence assay, a series of gradient dilution concentration of IS1081 plasmid from 8960 pM to 44.8 fM were prepared with 1 μL volume as RPA template. Meanwhile, no-MTB DNA was used as negative control and all of them were performed in three replicates. Total DNA mass for the assay was determined following formula: M/5-7×(4.38×10^6^×660) = μg/μL. M is concentration of IS1081 plasmid, 5-7 is number of copies of IS1081 per MTB genome and 4.38×10^6^ is approximately number of DNA length of M. bovis BCG. The specificity of the assay was obtained by analyzing at least 448 fM genomic DNA of target MTB, with/ or not closely related species from clinical isolates and standard strains.

### Clinical Performance Evaluation of MTB-Cas12a Fluorescence assay

The practical performance evaluation of the MTB detection assay was carried out via Cas12a trans-cleavage fluorescence system using a total of 194 clinical sputum samples, which were randomly collected from Fuzhou Center for Disease Control and Prevention in 2013~2016 with one sample came from one person, including patients with similar signs and symptoms consistent with pulmonary TB. All sample culture results, including 140 culture-positive results and 54 culture-negative results, were unknown for operator firstly. Genome of all samples were extracted using QIAGEN nucleic acid extraction Kit and the concentrations were measured via Nanodrop 2.0 and then stored at −20°C before test. The 194 clinical samples were detected by Cas12a/gRNA trans-cleavage fluorescence assay and the data analysis was carried out according to the above criteria.

### Data Analysis

All the data analysis was performed with SPSS version 2.2 and Origin Lab version 8.0, including ROC area, R square etc. Each experiment was repeated at 3 times for each sample.

## Results

### Design of Guide RNA of MTB

To ensure the coverage on all MTB species, the target insertion sequence IS1081 was selected for the assay and gRNAs targeting *mycobacterium tuberculosis* were designed. Principally, there are numbers of gRNAs corresponding to amounts of PAM structure existing in the insert sequence IS1081. Actually, different gRNAs had different target efficiency of Cas12a/gRNA system and not all predicted gRNAs were suitable for targeting MTB. Scores of target efficiency for gRNAs were evaluated by Cpf1-CRISPR-DT online software and range from 0.7061 to 0.97 was selected after excluding that of less than 0.6, and nine gRNAs were remained. In the end, we selected one that the score of target efficiency was 0.7061 based on unconspicuous secondary structure under reaction temperature and specificity of gRNA spacer for MTB. Especially, the spacer of MTB-specific sequence of gRNA was conserved region of IS1081 according to previous study and our sequence blasting. The sequences of designed IS1081 gRNAs and corresponding score of target efficiency for MTB genome were shown Table.2 in Supplemental Data.

**Table 2.** Clinical diagnostic performance of MTB-Cas12a assay on clinical samples

|  |  | MTB-Cas12a assay |  |  |
| --- | --- | --- | --- | --- |
|  |  | Positive | Negative | Total |
| Culture Method | Positive | 139 | 1 | 140 |
|  | Negative | 0 | 53 | 53 |
| Total |  | 139 | 54 | 193 |

### Feasibility Analysis of Purified Cas12a for MTB Detection

The gRNA-directed DNA binding and following activation of single-stranded DNA (ssDNA) trans-cleavage activity of Cas12a protein are vital properties for accurate recognition of target DNA for detection. The activity level of Cas12a protein directly determines the limit of detection of our Cas12a/gRNA trans-cleavage detection assay. As seen in Fig 2, when it exists IS1081 plasmid as template, the Cas12a/gRNA system, mixture of IS1081 gRNA and Cas12a protein, could degrade the target IS1081 plasmid *in cis* and single strand DNA *in trans*, resulting in reduction of the number of targets and non-target ssDNA (as reporter) in Cas12a/gRNA system or increase fluorescence intensity in Cas12a/gRNA fluorescence detection system. Excitingly, we observed IS1081 plasmid and ssDNA reporters were cleaved almost at the same time. These results suggested that the cis- and trans-cleavage activity ability of purified Cas12a protein is quite high and Cas12a/gRNA system for MTB detection is feasible.

**Figure 2.**
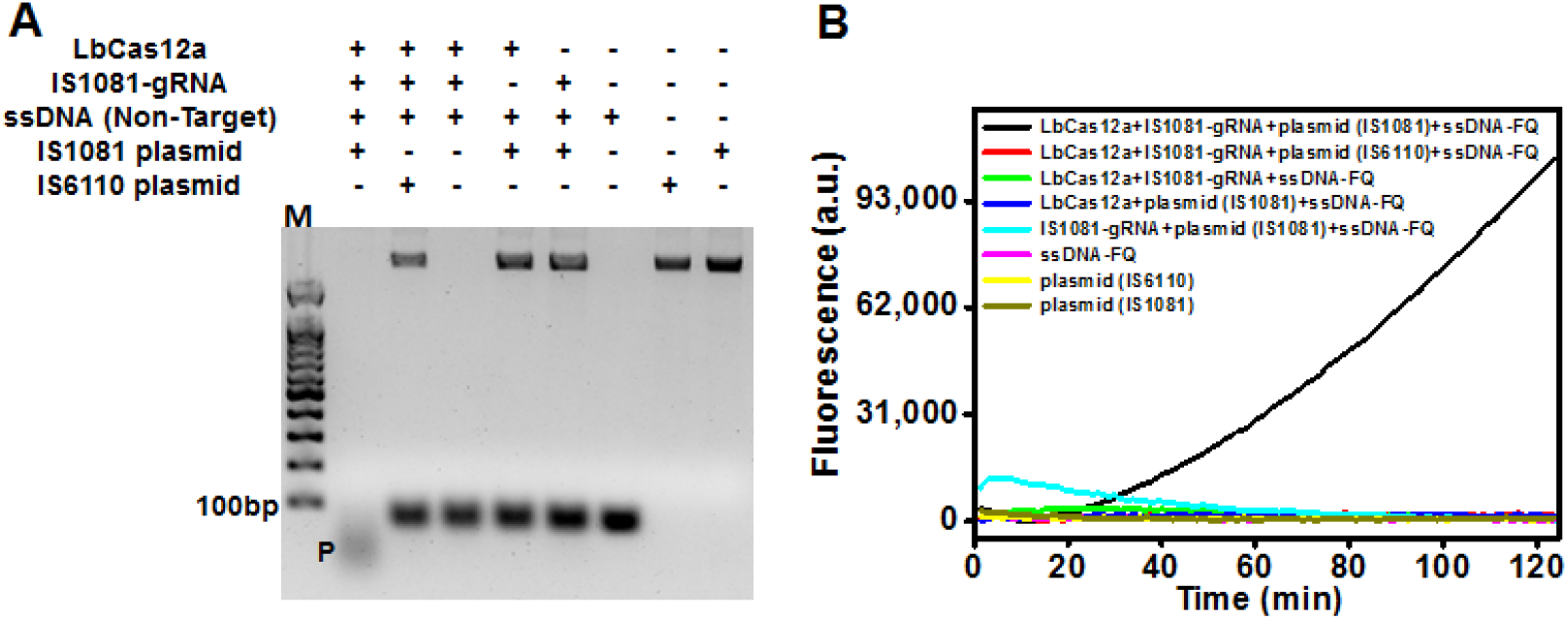
Feasibility analysis of Cas12a/gRNA system for MTB detection. 4% agarose gel electrophoretic analysis of the feasibility of Cas12a/gRNA system for MTB detection. 79 nt ssDNA (Non-Target) as substrate of Cas12a/gRNA trans cleavage (Figure 2A); Real-time fluorescence of Cas12a/gRNA fluorescence assay. ssDNA-FQ was served as the reporter probe. The concentration of the reaction components was as follows: 50nM purified LbCas12a, 36 nM IS1081-gRNA, 50 nM ssDNA (Non-Target) or ssDNA-FQ, 80 ng IS1081 or IS6110 plasmid (Figure 2B).

### Optimization of Cas12a Fluorescence Assay for MTB

In order to simplify operation to achieve “one-pot” detection pattern and improve reaction efficiency, the volume of RPA reaction was controlled under 12.5 μL and then added compositions of Cas12a/gRNA system to the final volume of 20 μL for Cas12a/gRNA trans-cleavage fluorescence assay. Prominently, the RPA reaction conditions were optimized to maximize amplification efficiency, including reaction temperature, concentration of dNTP and primers and Mg^2+^. As shown in Fig. 3 (details can be seen in Figure.2 in the Supplemental Data), we found 0.48 μM forward and reverse primers and 28 mM MgOAc of RPA reaction were mainly influence factors for Cas12a/gRNA trans-cleavage fluorescence assay, and the assay obtained higher fluorescence intensity under optimized conditions using the same concentration of target. Therefore, these conditions were used in the following experiments for MTB detection.

**Figure 3.**
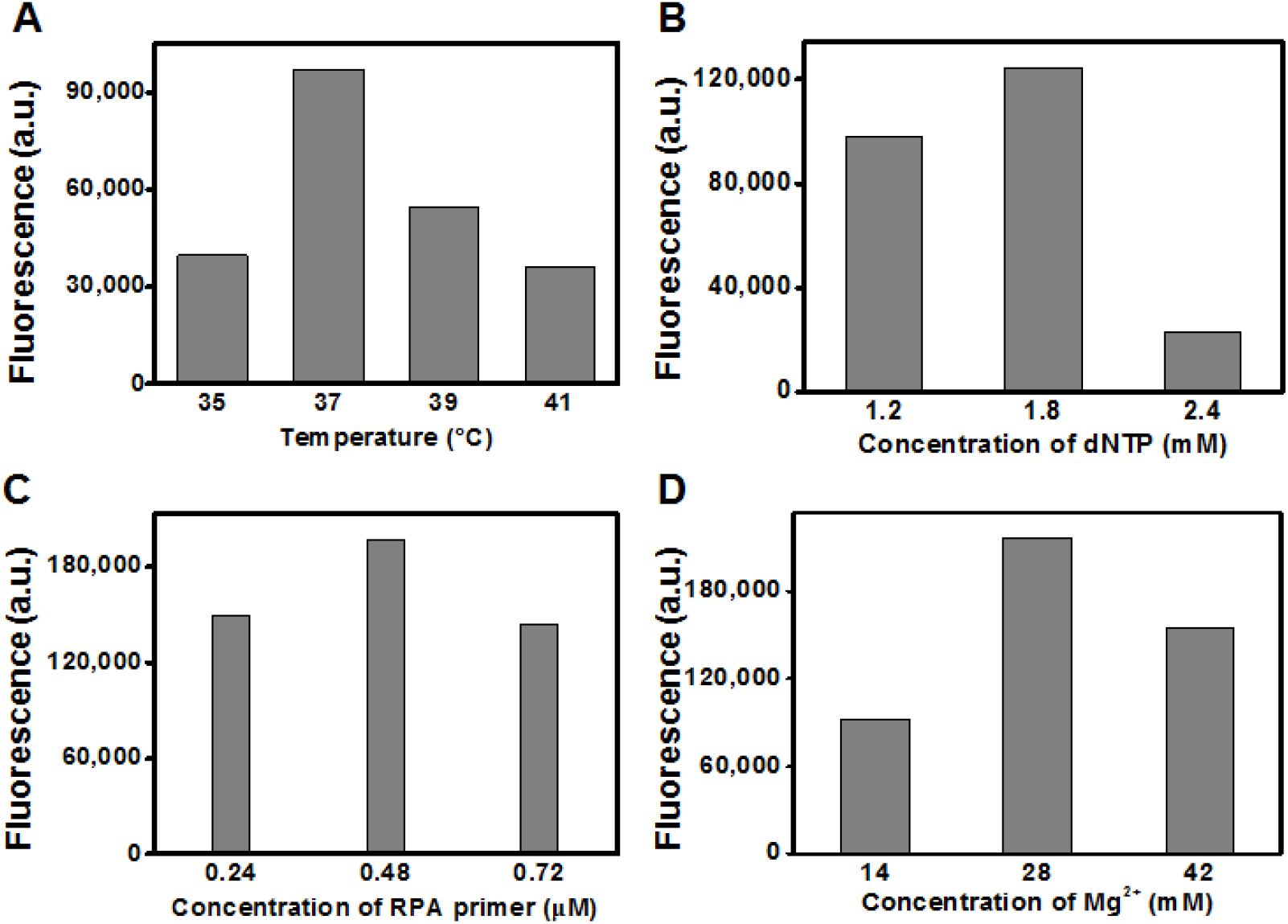
Optimization of MTBC-Cas12a trans-cleavage fluorescence assay. Figure 3A, Figure 3B, Figure 3C and Figure 3D are optimized results of assay on reaction temperature, dNTP concentration, primers concentration, and Mg^2+^ concentration, respectively.

### Performance of Cas12a Fluorescence Assay for MTB

To evaluate the ability of MTB-Cas12a trans-cleavage fluorescence assay, a series of gradient concentration of diluted IS1081 plasmids and genome of pathogens that may infect human respiratory tract were tested. For the sensitivity of MTB-Cas12a fluorescence assay, the fluorescence accumulation value of the assay decreased gradually with the decrease of concentration of IS1081 plasmid, and the limit of detection (LOD) for IS1081 was 4.48 fM (Fig. 4A). According to the formula depicted above, it was equivalent to 2.59-1.85 μg/L of genome of M. bovis BCG strain based on formula above from clinical samples. Comparing to RPA method^18^ for IS1081 of MTB detection with LOD of 20 fg for 50μL volume, it achieved four orders of magnitude higher and got attogram level which would dramatically promote positive detection rate. Meanwhile, the fluorescence value was proportional to the logarithm value of target concentrations over 6 orders of magnitude. The regression equation was Y=87983.47X+1.29307E6 with R square of 0.9775 (Fig. 4B). It was helpful to quantify MTB DNA accurately, especially regarding to low concentration of MTB DNA as substrate.

**Figure 4.**
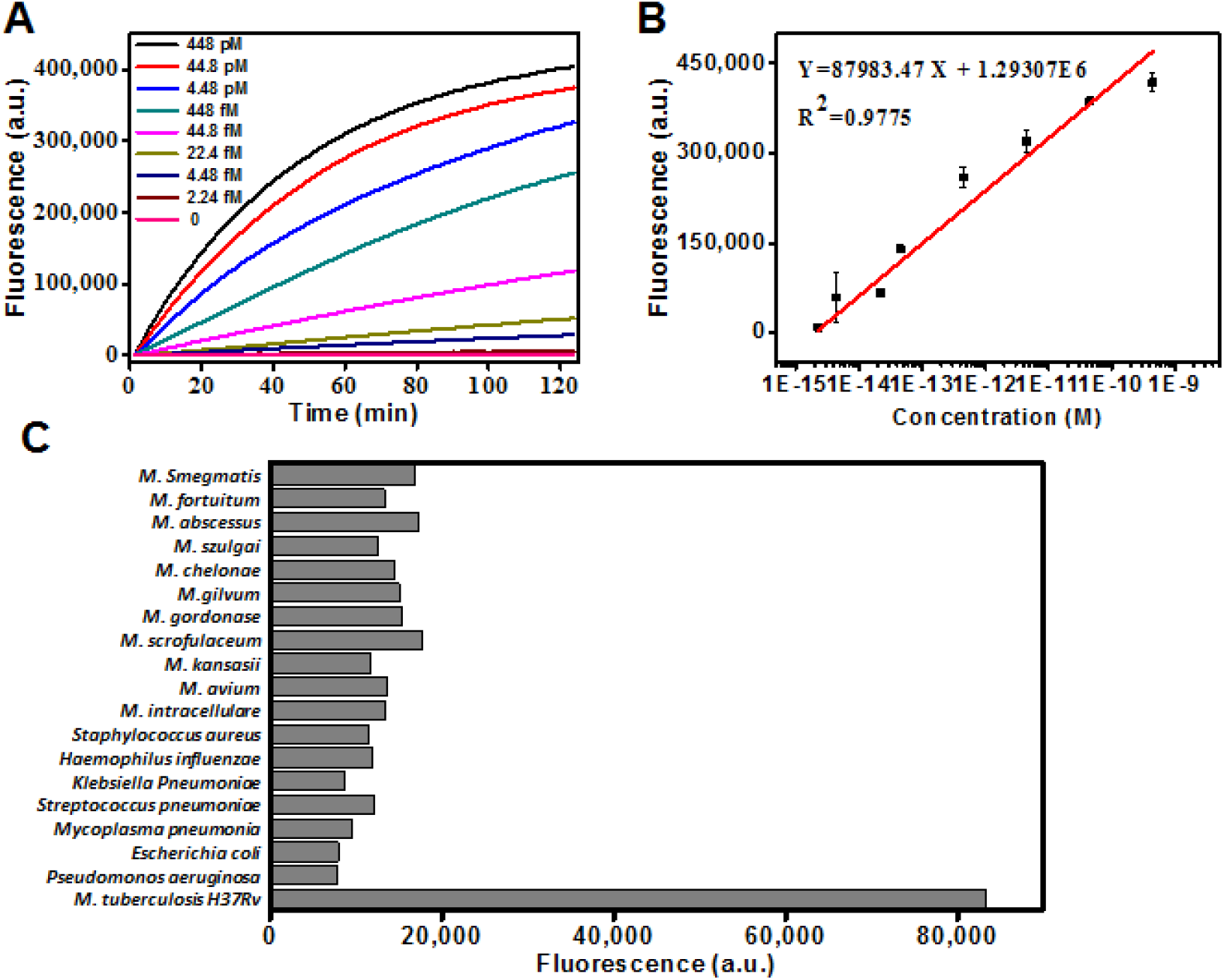
Detection performance of MTB-Cas12a fluorescence assay. Real-time fluorescence intensity alteration of MTB-Cas12a assay of different concentration of target(Figure 4A); The calibration plots of fluorescence intensity versus the logarithm of target concentration(Figure 4B); The specificity of the MTB-Cas12a fluorescence assay, including NTM and other pathogens of respiratory tract(Figure 4C).

To assess the specificity of the MTB-Cas12a fluorescence assay, genome of samples from a panel of NTM and pathogens which may infect human respiratory tract or their clinical symptoms similar to TB were tested. Three replicates were performed and 500 pg of DNA from either M. bovis BCG or M. tuberculosis H37Rv was used as positive template controls for each batch. Consequently, as shown in Fig. 4C, according to the judgment of criteria above, no positive results were obtained from the tested DNA except for the genome of MTB strains, indicating the high specificity of MTB-Cas12a fluorescence assay for MTB detection.

To evaluate the clinical diagnostic performance of the MTB-Cas12a assay, 193 patient samples (140 positive and 53 negative for MTB) which had been confirmed by the gold standard culture method were tested. Consequently, the fluorescence assay successfully detected 139 positive samples and 53 negative samples among 140 positive samples and 53 negative samples for MTB culture method. The detection accuracy of the assay based on 193 samples showed sensitivity of 99.29% (139/140) and specificity of 100% (53/53) (Table 2). Its performance was nearly similar to culture method except that one of culture-positive sample was failed to be detected whose fluorescence intensity was lower 1.5 times than that of negative result. Maybe this one failed to be detected by the assay due to the low concentration of genome and/or it was degraded in storage or during deliver process. Especially, the major reason could be that the MTB target for guide RNA recognition exist mutation resulting in none-complete hybridization, namely “turn-off” effect. As far as we can infer, it may be resolved by the modification of gRNA covering mutation site, such as LNA or leap over, similar to the processed methods of PCR and RPA for detection of mutation. But it needs to be proved in the future. In addition, Cas12a/gRNA trans cleavage fluorescence assay for MTB detection was required an average time of 4h, which including 1h of MTB DNA extraction, 1h for MTB DNA amplification by RPA and 2h for detection of Cas12a/gRNA system. It showed a significant advantage over culture method which needs few days, and a little slow compared to RPA method for MTB DNA detection which needs 1~2 h totally.

## Discussion

Rapid and sensitive detection method remains a challenge for clinical MTB diagnosis. Positive results of sputum smear microscopy were required at least five thousand of bacteria every milliliter and skilled technician. And culture method for MTB detection takes up to a few weeks and relatively low detection rate. They are not feasible for rapid molecular diagnostic of MTB in clinical applications. Rapid and sensitive MTB DNA detection method was attempted to be improved previously, such as automatic integration plate based PCR method—GeneXpert, RPA, etc. Unfortunately, they also brought some side effects which were similar to the original disadvantages and were not benefit for clinical MTB diagnosis. For example, continuous power input for temperature rising and falling and sample volume of 500 μL were required for GeneXpert. In contrast, Cas12a trans cleavage fluorescence assay for MTB can occur at 37°C which not needs a thermos-cycler and it could obtain positive results under low volume of MTB sample (100~200 μL). Meanwhile, it estimated to be 0.6138 dollars every sample^33^ comparing to 35~50 dollars for one sample of one cartridge using GeneXpert.

In this study, by optimizing the conditions of RPA reaction and Cas12a/gRNA system, we established one ultrasensitive method for Cas12a trans-cleavage fluorescence assay that could obtain 2.59-1.85 μg/L of limit of MTB genome (5-7 copies of IS1081 per MTB genome) detection from clinical samples. Although IS6110 is in multiple copies of up to 25 per genome of MTB and was used to be target in several commercial kits, such as Xpert MTB/RIF, the assay integrated the advantages of polymerase mediated DNA amplification and Cas12a mediated enzymatic signal amplification to compensate relatively low copies of IS1081 in 5-7 repeats per genome, and we proved that our Cas12a/gRNA system could detect MTB DNA with ultralow LOD. Also, Cas12a/gRNA system for IS1081 can strictly recognize MTB genome with PAM structure, which showed high specificity for target. Moreover, our study on 193 samples highlights the potential of Cas12a/gRNA as an ultra-sensitive, promising assay for diagnosis of clinical tuberculosis, in spite of miss detection of one sample of culture-positive for MTB which might due to the existence of mutation in the recognition sequence of Cas12-gRNA or low concentration of MTB DNA. So, in order to reduce the false negative possibility, spacer sequence of gRNA should be avoided from region with mutation sequences or SNP, especially mutation closes to PAM sequence.

In addition, there are also some limitations in our established Cas12a fluorescence assay. Firstly, it takes nearly twice time to get result compared with Xpert MTB\RIF. This may be reason the assay got ultra-sensitivity. But the time consumed totally could be controlled under the acceptable range for clinical diagnosis. Secondly, the Cas12a trans-cleavage fluorescence is detected by fluorescence detector but not visual readout that requires no additional devices. Finally, apart from diagnosis, drug resistance is also increasingly severe in the world for treating MTB patients. It is an urgent need to analyze mutations of drug susceptibility for one- or even second-line drugs via sensitive detection methods. With the natural characteristics of sensitive signal amplification of Cas/gRNA system, it has the potential to detect multiple mutations related to drug response. Ultra-sensitive and specific Cas12a/gRNA system combining polymerase mediated DNA amplification for both pathogen and drug resistance detection would permit the precise approach to control tuberculosis infection.

In a word, we have developed a CRISPR-Cas12a based fluorescence assay by combining isothermal recombinase polymerase amplification with Cas12a trans-cleavage activity, which could be activated by target specific DNA with high sensitivity and selectivity for rapid detection of pathogen *Mycobacterium tuberculosis* from clinical samples comparing to gold standard culture method which needs few days and extensive labor work. It is helpful to use this assay to prompt the confirmed diagnosis of mycobacterium tuberculosis complexes from none mycobacterium including NTM in clinical settings. But, in future, multi-center prospective study is needed to provide deeper understanding on its potential usage for clinical diagnosis.

## Abbreviations

TB: Tuberculosis
MTB: *mycobacterium tuberculosis*
CRISPR: Clustered Regularly Interspaced Short Palindromic Repeats
CRISPR-Cas: Clustered Regularly Interspaced Short Palindromic Repeats associated protein
gRNA: guide RNA
RPA: recombinase polymerase amplification
HIV: human immunodeficiency virus
AIDS: acquired immune deficiency syndrome
NTM: *non-tuberculosis mycobacteria*
PCR: polymerase chain reaction
WHO: World Health Organization
LAMP: loop mediated amplification
HDA: helicase dependent amplification
SNP: single nucleotide polymorphism
PAM: protospacer adjacent motif
LOD: limit of detection
ssDNA: single-stranded DNA

## Acknowledgments

This work was supported by the National Natural Science Foundation of China (Grant No. 21605021 and 21705022), the China Postdoctoral Science Foundation, the Scientific Foundation of Fujian Health Department (Grant No. 2019-1-87), the Scientific Foundation of Fuzhou Science and Technology Department (Grant No. 2019-S-90).

## References

1. World Health Organization (2018) Global Tuberculosis Report.

2. World Health Organization (2009) Global Tuberculosis Report.

3. Denkinger CM, Kik SV, Madhukar P Robust, reliable and resilient: designing molecular tuberculosis tests for microscopy centers in developing countries. Expert Review of Molecular Diagnostics 2013, 13:763–767.

4. Griffith DE, Timothy A, Brown-Elliott BA, Antonino C, Charles D, Fred G, Holland SM, Robert H, Gwen H, Iademarco MF An official ATS/IDSA statement: diagnosis, treatment, and prevention of nontuberculous mycobacterial diseases. Am J Respir Crit Care Med 2007, 175:367–416.

5. However, Pulmonary ATO Species Identification and Clarithromycin Susceptibility Testing of 278 Clinical Nontuberculosis Mycobacteria Isolates. Biomed Research International 2015, 2015:506598.

6. Wang X LH, Jiang G, Zhao L, Ma Y, Javid B, Huang H. Prevalence and drug resistance of nontuberculous mycobacteria, northern China, 2008-2011. Emerging Infect Dis 2014, 20:1252–1253.

7. Parrish NM, Carroll KC Role of the clinical mycobacteriology laboratory in diagnosis and management of tuberculosis in low-prevalence settings. Journal of Clinical Microbiology 2011, 49:772–776.

8. Sgaragli G, Frosini M Human Tuberculosis I. Epidemiology, Diagnosis and Pathogenetic Mechanisms. Current Medicinal Chemistry 2016, 23:2836–2873.

9. Machado Diana CI, Viveiros Miguel, Advances in the molecular diagnosis of tuberculosis: From probes to genomes. Infect Genet Evol 2019, 72:93–112.

10. World Health Organisation (2010) Strategic and Technical Advisory Group for Tuberculosis Report of 10th Meeting. Geneva.

11. Nimitphak T, Kiatpathomchai W, Flegel TW Loop-mediated isothermal amplification of DNA. Nucleic Acids Research 2000, 28:E63.

12. Vincent M, Xu Y, Kong H Helicase-dependent isothermal DNA amplification. Embo Reports 2004, 5:795–800.

13. Piepenburg O, Williams C, Stemple D, Na: DNA detection using recombination proteins. Plos Biology 2006, 4:1115–1121.

14. Li J, Macdonald J, Stetten FV Review: a comprehensive summary of a decade development of the recombinase polymerase amplification. Analyst 2018, 144:31–67.

15. Daher RK, Stewart G, Boissinot M, Bergeron MG Recombinase Polymerase Amplification for Diagnostic Applications. Clinical Chemistry 2016, 62:947–958.

16. Boehme CC, Pamela N, German H, Rubhana R, Zeaur R, Martina G, Erica S, Michael H, Tsugunori N, Tetsu H Operational feasibility of using loop-mediated isothermal amplification for diagnosis of pulmonary tuberculosis in microscopy centers of developing countries. Journal of Clinical Microbiology 2007, 45:1936–1940.

17. Wanyuan A, Stephen A, Evelyn W, Brian H, Larry R, Barry K, Robert J Rapid detection of rpoB gene mutations conferring rifampin resistance in Mycobacterium tuberculosis. Journal of Clinical Microbiology 2012, 50:2433–2440.

18. Boyle DS, Mcnerney R, Teng LH, Leader BT, Pérez-Osorio AC, Meyer JC, O’Sullivan DM, Brooks DG, Piepenburg O, Forrest MS Rapid detection of Mycobacterium tuberculosis by recombinase polymerase amplification. PLoS One 2014, 9:e103091.

19. Daher RK, Stewart G, Boissinot M, Boudreau DK, Bergeron MG Influence of sequence mismatches on the specificity of recombinase polymerase amplification technology. Molecular & Cellular Probes 2015, 29:116–121.

20. Nguyen DTT, Lee EY, Koo B, Jin CE, Lee TY, Yong S A microfluidic enrichment platform with a recombinase polymerase amplification sensor for pathogen diagnosis. Analytical Biochemistry 2017, 544:87–92.

21. Peng Z, Gao W, Huang H, Jiang J, Chen X, Fan J, Yan X Rapid Detection of Vibrio parahaemolyticus in Shellfish by Real-Time Recombinase Polymerase Amplification. Food Analytical Methods 2018, 11:1–9.

22. Kersting S, Rausch V, Bier FF, Nickisch-Rosenegk MV Multiplex isothermal solid-phase recombinase polymerase amplification for the specific and fast DNA-based detection of three bacterial pathogens. Microchimica Acta 2014, 181:1715–1723.

23. Mauk MG, Liu C, Sadik M, Bau HH Microfluidic Devices for Nucleic Acid (NA) Isolation, Isothermal NA Amplification, and Real-Time Detection. Methods in Molecular Biology 2015, 1256:15.

24. Yamanaka ES, Tortajada-Genaro LA, Maquieira Á Low-cost genotyping method based on allele-specific recombinase polymerase amplification and colorimetric microarray detection. Microchimica Acta 2017, 184:1453–1462.

25. Liu X YQ, Huang J, Chen J, Guo Z, Liu Z, Cai L, Li R, Wang Y, Yang G, Lan Q. Influence of design probe and sequence mismatches on the efficiency of fluorescent RPA. World Journal of Microbiology and Biotechnology 2019, 35:95.

26. Chen JS, Ma E, Harrington LB, Costa MD, Doudna JA CRISPR-Cas12a target binding unleashes indiscriminate single-stranded DNase activity. Science 2018, 360:436–439.

27. Martin J, Krzysztof C, Ines F, Michael H, Doudna JA, Emmanuelle C A programmable dual-RNA-guided DNA endonuclease in adaptive bacterial immunity. Science 2012, 337:816–821.

28. Rodolphe B, Christophe F, Hélène D, Melissa R, Patrick B, Sylvain M, Romero DA, Philippe H CRISPR provides acquired resistance against viruses in prokaryotes. Science 2007, 315:1709–1712.

29. Ran FA, Hsu PD, Wright J, Agarwala V, Scott DA, Zhang F Genome engineering using the CRISPR-Cas9 system. Nature Protocols 2013, 8:2281.

30. Le C, F Ann R, David C, Shuailiang L, Robert B, Naomi H, Hsu PD, Xuebing W, Wenyan J, Marraffini LA Multiplex genome engineering using CRISPR/Cas systems. Science 2015, 339:819–823.

31. Manguso RT, Pope HW, Zimmer MD, Brown FD, Yates KB, Miller BC, Collins NB, Bi K, Lafleur MW, Juneja VR In vivo CRISPR screening identifies Ptpn2 as a cancer immunotherapy target. Nature 2017, 547:413.

32. Lu XJ, Xue HY, Ke ZP, Chen JL, Ji LJ CRISPR-Cas9: a new and promising player in gene therapy. Journal of Medical Genetics 2015, 52:289–296.

33. Gootenberg JS AO, Lee JW, Essletzbichler P, Dy AJ, Joung J, Verdine V, Donghia N, Daringer NM, Freije CA, Myhrvold C, Bhattacharyya RP, Livny J, Regev A, Koonin EV, Hung DT, Sabeti PC, Collins JJ, Zhang F. Nucleic acid detection with CRISPR-Cas13a/C2c2. Science 2017, 356:438–442.

34. Gootenberg JS, Abudayyeh OO, Kellner MJ, Joung J, Collins JJ, Zhang F Multiplexed and portable nucleic acid detection platform with Cas13, Cas12a, and Csm6. Science 2018, 360:439–444.

35. Harrington LB, Burstein D, Chen JS, Paez-Espino D, Ma E, Witte IP, Cofsky JC, Kyrpides NC, Banfield JF, Doudna JA Programmed DNA destruction by miniature CRISPR-Cas14 enzymes. Science 2018, 362:839–842.

36. Collins D M SDM Identification of an insertion sequence, IS1081, in Mycobacterium bovis. FEMS Microbiology Letters 1991, 83:11–16.

37. Ma Q, Liu H, Ye F, Xiang G, Shan W, Xing W Rapid and visual detection of Mycobacterium tuberculosis complex using recombinase polymerase amplification combined with lateral flow strips. Molecular & Cellular Probes 2017, 36:43–49.

