## Supplementary materials for "An isothermal method for sensitive detection of *Mycobacterium tuberculosis complexes* using CRISPR/Cas12a *cis-* and *trans-*cleavage"

**Supplementary material**

**Supplemental Table 1. The oligonucleotide sequences of RPA primers for selection**

| RPA primer | Sequence ($\boldsymbol{5'\to3'}$) |
| --- | --- |
| IS1081-F1 | CCAAGCTGCGCCAGGGCAGCTATTTCCCGGAC |
| IS1081-F2 | CAAGCGAGCTGAACGCGCACTGACCAGCGT |
| IS1081-F3 | GGTGGCGACCTGCTACCTGCTGGGAGTAT |
| IS1081-R1 | TTGGCCATGATCGACACTTGCGACTTGGA |
| IS1081-R2 | CGAAACGCCTCTACGGCTTCGTCGAGCTC |
| IS1081-R3 | GGCGTCGGCGGCGAGGAAGGTATACGG |

**Supplemental Table 2. The sequences of designed IS1081 gRNAs and corresponding score of target efficiency for MTB genome**

| **Number** | **Sequence (**$\boldsymbol{5'\to3'}$**)** | **Score of target efficiency** |
| --- | --- | --- |
| **1** | UAAUUUCUACUAAGUGUAGAUGUCAACCCAGCACCUGCCAG | 0.9725 |
| **2** | UAAUUUCUACUAAGUGUAGAUGACACCCGTGCCGCAACCAT | 0.9639 |
| **3** | UAAUUUCUACUAAGUGUAGAUGAAGGAUCACGCGAUGACCG | 0.8954 |
| **4** | UAAUUUCUACUAAGUGUAGAUACCACACCUUGGGGCACCUU | 0.8461 |
| **5** | UAAUUUCUACUAAGUGUAGAUGCCAUGAUCGACACUUGCGA | 0.838 |
| **6** | UAAUUUCUACUAAGUGUAGAUCAAGUCGCAAGUGUCGAUCA | 0.7736 |
| **7** | UAAUUUCUACUAAGUGUAGAUUCGGUCAGAGCGUCGAGUAC | 0.7692 |
| **8** | UAAUUUCUACUAAGUGUAGAUGGACCCGCCCGCUCGAUGCC | 0.7227 |
| **9** | UAAUUUCUACUAAGUGUAGAUGACCAGGCGCUCCAUCCGGC | 0.7061 |
| **10** | UAAUUUCUACUAAGUGUAGAUGCGCCAGAUCUGCUUGGGGA | 0.5644 |
| **11** | UAAUUUCUACUAAGUGUAGAUUCACACCAAGUGUUUCGACC | 0.5353 |
| **12** | UAAUUUCUACUAAGUGUAGAUCUGGUCGUUUCGAAGGAUCA | 0.4128 |
| **13** | UAAUUUCUACUAAGUGUAGAUCCGGACUGGCUGCUGCAGCG | 0.2013 |
| **14** | UAAUUUCUACUAAGUGUAGAUCUGCUUGGCGGGUUCUUCGG | 0.0606 |
| **15** | UAAUUUCUACUAAGUGUAGAUUUUCCUUCGAUUGACUCUGG | 0.0107 |

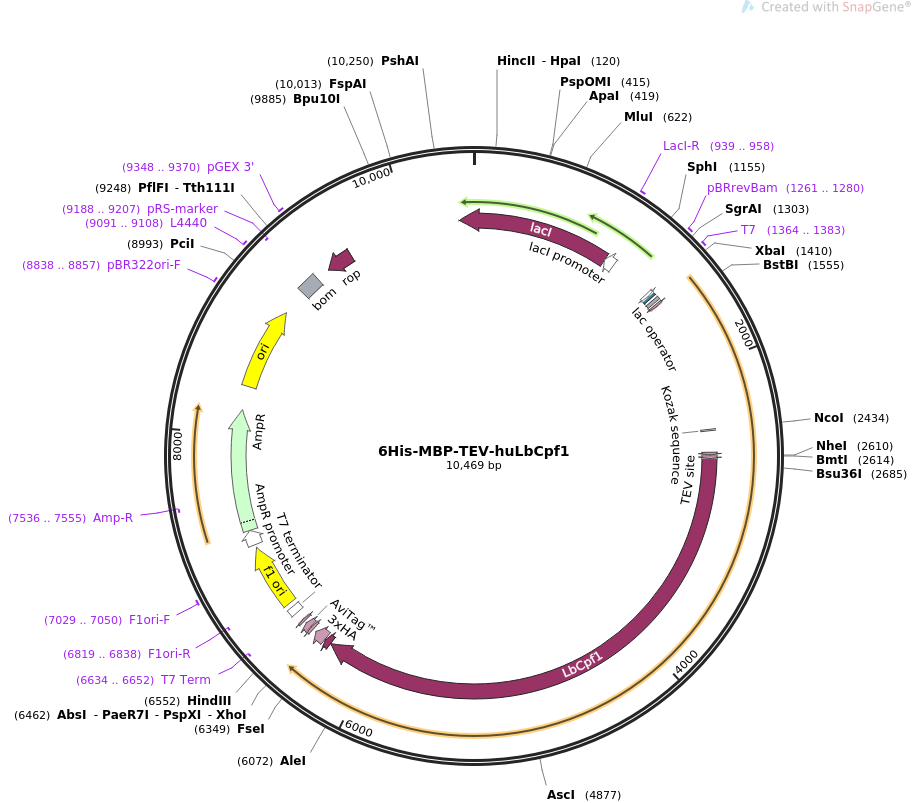

**Supplemental Figure 1. Map of plasmid of LbCas12a protein expression**

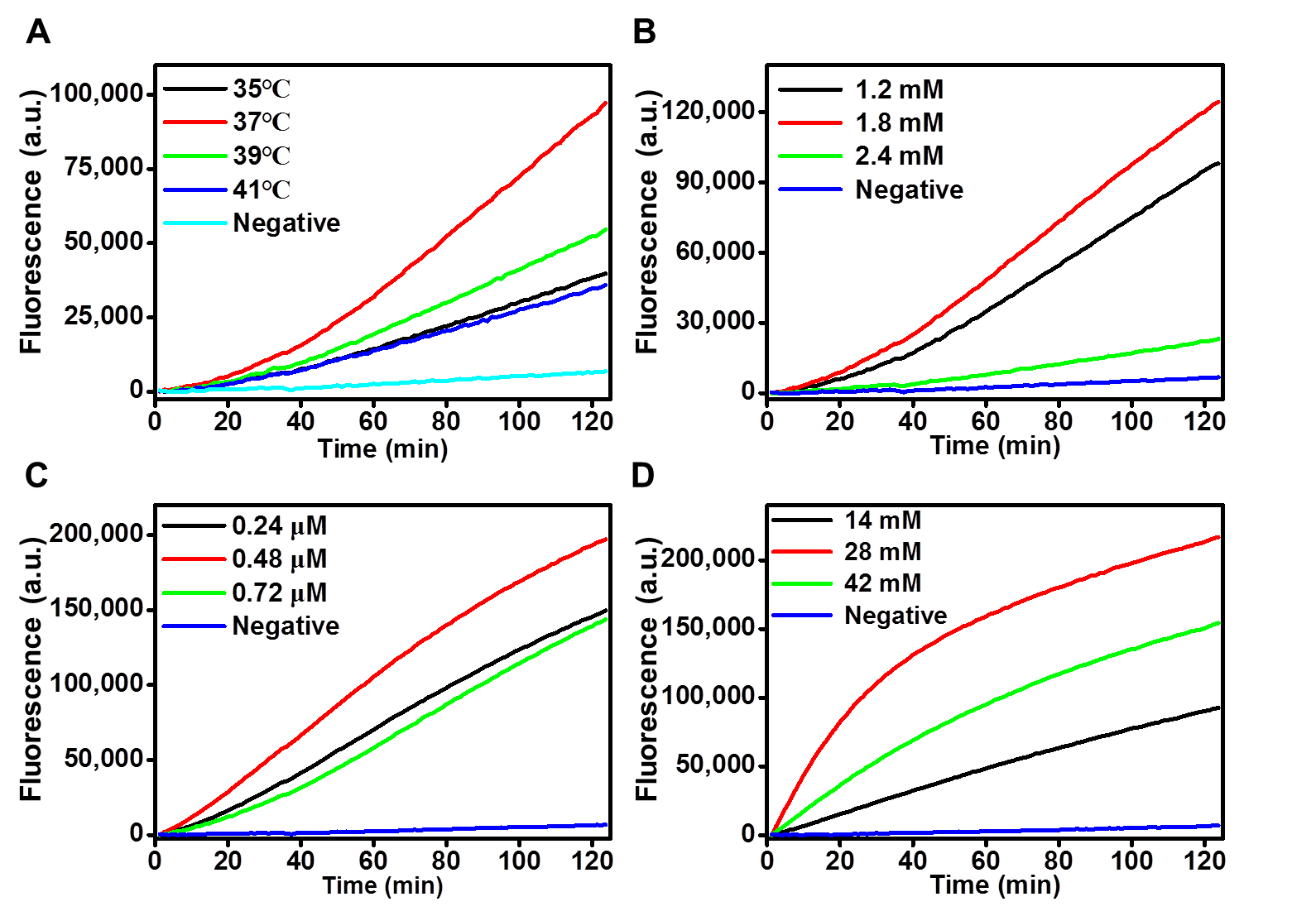

**Supplemental Figure 2. Optimization of MTBC-Cas12a trans-cleavage fluorescence assay** A, B, C, D are optimized fluorescence intensity & time of the assay on reaction temperature, dNTP concentration, primers concentration, and Mg^2+^ concentration, respectively.

**Supplementary Information for IS6110\ IS1081\ LbCas12a protein expression plasmid**

**Supplementary M1. Mycobacterium tuberculosis IS6110 insertion sequence (Genbank ID: X17348.1) cloned to PUC57 vector**

CGATGAACCGCCCCGGCATGTCCGGAGACTCCAGTTCTTGGAAAGGATGGGGTCATGTCAGGTGGTTCATCGAGGAGGTACCCGCCGGAGCTGCGTGAGCGGGCGGTGCGGATGGTCGCAGAGATCCGCGGTCAGCACGATTCGGAGTGGGCAGCGATCAGTGAGGTCGCCCGTCTACTTGGTGTTGGCTGCGCGGAGACGGTGCGTAAGTGGGTGCGCCAGGCGCAGGTCGATGCCGGCGCACGGCCCGGGACCACGACCGAAGAATCCGCTGAGCTGAAGCGCTTAGCGGCGGGACAACGCCGAATTGCGAAGGGCGAACGCGATTTTAAAGACCGCGTCGGCTTTCTTCGCGGCCGAGCTCGACCGGCCAGCACGCTAATTAACGGTTCATCGCCGATCATCAGGGCCACCGCGAGGGCCCCGATGGTTTGCGGTGGGGTGTCGAGTCGATCTGCACACAGCTGACCGAGCTGGGTGTGCCGATCGCCCCATCGACCTACTACGACCACATCAACCGGGAGCCCAGCCGCCGCGAGCTGCGCGATGGCGAACTCAAGGAGCACATCAGCCGCGTCCACGCCGCCAACTACGGTGTTTACGGTGCCCGCAAAGTGTGGCTAACCCTGAACCGTCAGGGCATCGAGGTGGCCAGATGCACCGTCGAACGGCTGATGACCAAACTCGGCCTGTCCGGGACCACCCGCGGCAAAGCCCGCAGGACCACGATCGCTGATCCGGCCACAGCCCGTCCCGCCGATCTCGTCCAGCGCCGCTTCGGACCACCAGCACCTAACCGGCTGTGGGTAGCAGACCTCACCTATGTGTCGACCTGGGCAGGGTTCGCCTACGTGGCCTTTGTCACCGACGCCTACGCTCGCAGGATCCTGGGCTGGCGGGTCGCTTCCACGATGGCCACCTCCATGGTCCTCGACGCGATCGAGCAAGCCATCTGGACCCGCCAACAAGAAGGCGTACTCGACCTGAAAGACGTTATCCACCATACGGATAGGGGATCTCAGTACACATCGATCCGGTTCAGCGAGCGGCTCGCCGAGGCAGGCATCCAACCGTCGGTCGGAGCGGTCGGAAGCTCCTATGACAATGCACTAGCCGAGACGATCAACGGCCTATACAAGACCGAGCTGATCAAACCCGGCAAGCCCTGGCGGTCCATCGAGGATGTCGAGTTGGCCACCGCGCGCTGGGTCGACTGGTTCAACCATCGCCGCCTCTACCAGTACTGCGGCGACGTCCCGCCGGTCGAACTCGAGGCTGCCTACTACGCTCAACGCCAGAGACCAGCCGCCGGCTGAGGTCTCAGATCAGAGAGTCTCCGGACTCACCGGGGCGGTTCACGA

**Supplementary M2. Mycobacterium tuberculosis IS1081 insertion sequence (Genbank ID: CP003248.2) cloned to PUC57 vector**

GAATTCGATCGCCGAGCCGACAAGACATGCCAGCGCAACCCGCTTCATCGTCGTGGCAGGTGTTGGGCTGATTTTGGTCAACCCAGCACCTGCCAGGACGGGCTACGGATGTACACGGCGACGACGGTATGGGAGGATGTCCGGTCTTGCTCCGGTCATGTCCGGTGAATGTGCTGCCAACATCCTGGGGACCGTCCAGCGAGTTTCACCACACCTTGGGGCACCTTCTGTCACTGCTCGGTGCTGTGGATTGGTGTCAAGTTACGTCCAGGGGTGTGGTGTACGGGCAGGTAAGGCCGGTGGGCGTGTCGTAGCCCAGTAGTGGGCGGTCATCGCGTGATCCTTCGAAACGACCAGCAAAAGTCAATCGAAGGAAATGACGCAATGACCTCTTCTCATCTTATCGACACCGAGCAGCTTCTGGCTGACCAACTCGCACAGGCGAGCCCGGATCTGCTGCGCGGGCTGCTCTCGACGTTCATCGCCGCCTTGATGGGGGCTGAAGCCGACGCCCTGTGCGGGGCGGGCTACCGCGAACGCAGCGATGAGCGGTCCAATCAGCGCAACGGCTACCGCCACCGTGATTTCGACACCCGTGCCGCAACCATCGACGTCGCGATCCCCAAGCTGCGCCAGGGCAGCTATTTCCCGGACTGGCTGCTGCAGCGCCGCAAGCGAGCTGAACGCGCACTGACCAGCGTGGTGGCGACCTGCTACCTGCTGGGAGTATCCACTCGCCGGATGGAGCGCCTGGTCGAAACACTTGGTGTGACAAAGCTTTCCAAGTCGCAAGTGTCGATCATGGCCAAAGAGCTCGACGAAGCCGTAGAGGCGTTTCGGACCCGCCCGCTCGATGCCGGCCCGTATACCTTCCTCGCCGCCGACGCCCTGGTGCTCAAGGTGCGCGAGGCAGGCCGCGTCGTCGGAGTGCACACCTTGATCGCCACCGGCGTCAACGCCGAGGGCTACCGAGAGATCCTGGGCATCCAGGTCACCTCCGCCGAGGACGGGGCCGGCTGGCTGGCGTTCTTCCGCGACCTGGTCGCCCGCGGCCTGTCCGGGGTCGCGCTGGTCACCAGCGACGCCCACGCCGGCCTGGTGGCCGCGATCGGCGCCACCCTGCCCGCAGCGGCCTGGCAGCGCTGCAGAACCCACTACGCAGCCAATCTGATGGCAGCCACCCCGAAGCCCTCCTGGCCGTGGGTGCGCACCCTGCTGCACTCCATCTACGACCAGCCCGACGCCGAATCAGTTGTTGCCCAATATGATCGGGTACTCGACGCTCTGACCGACAAACTCCCCGCGGTGGCCGAGCACCTCGACACCGCCCGCACCGACCTGCTGGCGTTCACCGCCTTCCCCAAGCAGATCTGGCGCCAAATCTGGTCCAACAACCCCCAGGAACGCCTCAACCGAGAGGTACGACGCCGAACCGACGTCGTGGGCATCTTCCCCGACCGCGCCTCGATCATCCGCCTCGTCGGAGCCGTCCTCGCCGAACAACACGACGAATGGATCGAAGGACGGCGCTACCTGGGCCTCGAGGTCCTCACCCGAGCCCGAGCAGCACTGACCAGCACCGAAGAACCCGCCAAGCAGCAAACCACCAACACCCCAGCACTGACCACCTAGACTGCCACCCGAAGGATCACGCGAGGAACCTTCACTCGTACACCACGTCCCTGGCCTTGGCCTGGTGTCAGGCCCAGCTG

**Supplementary M3. Plasmid sequence of LbCas12a protein expression**

CTTTTCACCAGTGAGACGGGCAACAGCTGATTGCCCTTCACCGCCTGGCCCTGAGAGAGTTGCAGCAAGCGGTCCACGCTGGTTTGCCCCAGCAGGCGAAAATCCTGTTTGATGGTGGTTAACGGCGGGATATAACATGAGCTGTCTTCGGTATCGTCGTATCCCACTACCGAGATATCCGCACCAACGCGCAGCCCGGACTCGGTAATGGCGCGCATTGCGCCCAGCGCCATCTGATCGTTGGCAACCAGCATCGCAGTGGGAACGATGCCCTCATTCAGCATTTGCATGGTTTGTTGAAAACCGGACATGGCACTCCAGTCGCCTTCCCGTTCCGCTATCGGCTGAATTTGATTGCGAGTGAGATATTTATGCCAGCCAGCCAGACGCAGACGCGCCGAGACAGAACTTAATGGGCCCGCTAACAGCGCGATTTGCTGGTGACCCAATGCGACCAGATGCTCCACGCCCAGTCGCGTACCGTCTTCATGGGAGAAAATAATACTGTTGATGGGTGTCTGGTCAGAGACATCAAGAAATAACGCCGGAACATTAGTGCAGGCAGCTTCCACAGCAATGGCATCCTGGTCATCCAGCGGATAGTTAATGATCAGCCCACTGACGCGTTGCGCGAGAAGATTGTGCACCGCCGCTTTACAGGCTTCGACGCCGCTTCGTTCTACCATCGACACCACCACGCTGGCACCCAGTTGATCGGCGCGAGATTTAATCGCCGCGACAATTTGCGACGGCGCGTGCAGGGCCAGACTGGAGGTGGCAACGCCAATCAGCAACGACTGTTTGCCCGCCAGTTGTTGTGCCACGCGGTTGGGAATGTAATTCAGCTCCGCCATCGCCGCTTCCACTTTTTCCCGCGTTTTCGCAGAAACGTGGCTGGCCTGGTTCACCACGCGGGAAACGGTCTGATAAGAGACACCGGCATACTCTGCGACATCGTATAACGTTACTGGTTTCACATTCACCACCCTGAATTGACTCTCTTCCGGGCGCTATCATGCCATACCGCGAAAGGTTTTGCGCCATTCGATGGTGTCCGGGATCTCGACGCTCTCCCTTATGCGACTCCTGCATTAGGAAGCAGCCCAGTAGTAGGTTGAGGCCGTTGAGCACCGCCGCCGCAAGGAATGGTGCATGCAAGGAGATGGCGCCCAACAGTCCCCCGGCCACGGGGCCTGCCACCATACCCACGCCGAAACAAGCGCTCATGAGCCCGAAGTGGCGAGCCCGATCTTCCCCATCGGTGATGTCGGCGATATAGGCGCCAGCAACCGCACCTGTGGCGCCGGTGATGCCGGCCACGATGCGTCCGGCGTAGAGGATCGAGATCTCGATCCCGCGAAATTAATACGACTCACTATAGGGGAATTGTGAGCGGATAACAATTCCCCTCTAGAAATAATTTTGTTTAACTTTAAGAAGGAGATATACATATGGGTCATCATCATCACCATCACAAAATCGAAGAAGGCAAACTGGTTATTTGGATCAATGGCGATAAAGGCTATAATGGTCTGGCAGAAGTTGGCAAAAAATTCGAAAAAGATACCGGCATTAAAGTGACCGTTGAACATCCGGATAAACTGGAAGAAAAATTTCCGCAGGTTGCAGCAACCGGTGATGGTCCGGATATTATCTTTTGGGCACATGATCGTTTTGGTGGTTATGCACAGAGCGGTCTGCTGGCAGAAATTACACCGGATAAAGCATTTCAGGACAAACTGTATCCGTTTACCTGGGATGCAGTTCGCTATAACGGTAAACTGATTGCATATCCGATTGCAGTTGAAGCACTGAGCCTGATCTATAACAAAGATCTGCTGCCGAATCCGCCTAAAACCTGGGAAGAAATTCCGGCACTGGATAAAGAACTGAAAGCAAAAGGTAAAAGCGCACTGATGTTTAATCTGCAAGAACCGTATTTTACCTGGCCTCTGATTGCAGCAGATGGTGGCTATGCATTCAAATATGAAAACGGCAAATATGATATCAAAGACGTGGGTGTTGATAATGCGGGTGCAAAAGCCGGTCTGACCTTTCTGGTTGATCTGATTAAAAACAAACACATGAACGCCGATACCGATTATAGCATTGCAGAAGCAGCATTTAACAAAGGTGAAACCGCAATGACAATTAATGGTCCGTGGGCATGGTCAAATATTGATACCAGCAAAGTGAATTATGGTGTTACCGTTCTGCCGACATTTAAAGGTCAGCCGAGCAAACCGTTTGTTGGTGTTCTGAGCGCAGGTATTAATGCAGCAAGCCCGAACAAAGAACTGGCAAAAGAATTTCTGGAAAACTATCTGCTGACCGATGAAGGTCTGGAAGCAGTGAATAAAGATAAACCGCTGGGTGCAGTTGCACTGAAAAGCTATGAAGAAGAACTGGTTAAAGATCCGCGTATTGCAGCCACCATGGAAAATGCACAGAAAGGCGAAATTATGCCGAATATTCCGCAGATGAGCGCATTTTGGTATGCCGTTCGTACCGCAGTGATTAATGCCGCATCAGGTCGTCAGACCGTTGATGAAGCCCTGAAAGATGCCCAGACCGGTGCAAGCACCGGATCCCTGGTTCCGCGTGGTAGCGCTAGCGAAAATCTGTATTTTCAGGGAATGAGCAAGCTGGAGAAGTTTACAAACTGCTACTCCCTGTCTAAGACCCTGAGGTTCAAGGCCATCCCTGTGGGCAAGACCCAGGAGAACATCGACAATAAGCGGCTGCTGGTGGAGGACGAGAAGAGAGCCGAGGATTATAAGGGCGTGAAGAAGCTGCTGGATCGCTACTATCTGTCTTTTATCAACGACGTGCTGCACAGCATCAAGCTGAAGAATCTGAACAATTACATCAGCCTGTTCCGGAAGAAAACCAGAACCGAGAAGGAGAATAAGGAGCTGGAGAACCTGGAGATCAATCTGCGGAAGGAGATCGCCAAGGCCTTCAAGGGCAACGAGGGCTACAAGTCCCTGTTTAAGAAGGATATCATCGAGACAATCCTGCCAGAGTTCCTGGACGATAAGGACGAGATCGCCCTGGTGAACAGCTTCAATGGCTTTACCACAGCCTTCACCGGCTTCTTTGATAACAGAGAGAATATGTTTTCCGAGGAGGCCAAGAGCACATCCATCGCCTTCAGGTGTATCAACGAGAATCTGACCCGCTACATCTCTAATATGGACATCTTCGAGAAGGTGGACGCCATCTTTGATAAGCACGAGGTGCAGGAGATCAAGGAGAAGATCCTGAACAGCGACTATGATGTGGAGGATTTCTTTGAGGGCGAGTTCTTTAACTTTGTGCTGACACAGGAGGGCATCGACGTGTATAACGCCATCATCGGCGGCTTCGTGACCGAGAGCGGCGAGAAGATCAAGGGCCTGAACGAGTACATCAACCTGTATAATCAGAAAACCAAGCAGAAGCTGCCTAAGTTTAAGCCACTGTATAAGCAGGTGCTGAGCGATCGGGAGTCTCTGAGCTTCTACGGCGAGGGCTATACATCCGATGAGGAGGTGCTGGAGGTGTTTAGAAACACCCTGAACAAGAACAGCGAGATCTTCAGCTCCATCAAGAAGCTGGAGAAGCTGTTCAAGAATTTTGACGAGTACTCTAGCGCCGGCATCTTTGTGAAGAACGGCCCCGCCATCAGCACAATCTCCAAGGATATCTTCGGCGAGTGGAACGTGATCCGGGACAAGTGGAATGCCGAGTATGACGATATCCACCTGAAGAAGAAGGCCGTGGTGACCGAGAAGTACGAGGACGATCGGAGAAAGTCCTTCAAGAAGATCGGCTCCTTTTCTCTGGAGCAGCTGCAGGAGTACGCCGACGCCGATCTGTCTGTGGTGGAGAAGCTGAAGGAGATCATCATCCAGAAGGTGGATGAGATCTACAAGGTGTATGGCTCCTCTGAGAAGCTGTTCGACGCCGATTTTGTGCTGGAGAAGAGCCTGAAGAAGAACGACGCCGTGGTGGCCATCATGAAGGACCTGCTGGATTCTGTGAAGAGCTTCGAGAATTACATCAAGGCCTTCTTTGGCGAGGGCAAGGAGACAAACAGGGACGAGTCCTTCTATGGCGATTTTGTGCTGGCCTACGACATCCTGCTGAAGGTGGACCACATCTACGATGCCATCCGCAATTATGTGACCCAGAAGCCCTACTCTAAGGATAAGTTCAAGCTGTATTTTCAGAACCCTCAGTTCATGGGCGGCTGGGACAAGGATAAGGAGACAGACTATCGGGCCACCATCCTGAGATACGGCTCCAAGTACTATCTGGCCATCATGGATAAGAAGTACGCCAAGTGCCTGCAGAAGATCGACAAGGACGATGTGAACGGCAATTACGAGAAGATCAACTATAAGCTGCTGCCCGGCCCTAATAAGATGCTGCCAAAGGTGTTCTTTTCTAAGAAGTGGATGGCCTACTATAACCCCAGCGAGGACATCCAGAAGATCTACAAGAATGGCACATTCAAGAAGGGCGATATGTTTAACCTGAATGACTGTCACAAGCTGATCGACTTCTTTAAGGATAGCATCTCCCGGTATCCAAAGTGGTCCAATGCCTACGATTTCAACTTTTCTGAGACAGAGAAGTATAAGGACATCGCCGGCTTTTACAGAGAGGTGGAGGAGCAGGGCTATAAGGTGAGCTTCGAGTCTGCCAGCAAGAAGGAGGTGGATAAGCTGGTGGAGGAGGGCAAGCTGTATATGTTCCAGATCTATAACAAGGACTTTTCCGATAAGTCTCACGGCACACCCAATCTGCACACCATGTACTTCAAGCTGCTGTTTGACGAGAACAATCACGGACAGATCAGGCTGAGCGGAGGAGCAGAGCTGTTCATGAGGCGCGCCTCCCTGAAGAAGGAGGAGCTGGTGGTGCACCCAGCCAACTCCCCTATCGCCAACAAGAATCCAGATAATCCCAAGAAAACCACAACCCTGTCCTACGACGTGTATAAGGATAAGAGGTTTTCTGAGGACCAGTACGAGCTGCACATCCCAATCGCCATCAATAAGTGCCCCAAGAACATCTTCAAGATCAATACAGAGGTGCGCGTGCTGCTGAAGCACGACGATAACCCCTATGTGATCGGCATCGATAGGGGCGAGCGCAATCTGCTGTATATCGTGGTGGTGGACGGCAAGGGCAACATCGTGGAGCAGTATTCCCTGAACGAGATCATCAACAACTTCAACGGCATCAGGATCAAGACAGATTACCACTCTCTGCTGGACAAGAAGGAGAAGGAGAGGTTCGAGGCCCGCCAGAACTGGACCTCCATCGAGAATATCAAGGAGCTGAAGGCCGGCTATATCTCTCAGGTGGTGCACAAGATCTGCGAGCTGGTGGAGAAGTACGATGCCGTGATCGCCCTGGAGGACCTGAACTCTGGCTTTAAGAATAGCCGCGTGAAGGTGGAGAAGCAGGTGTATCAGAAGTTCGAGAAGATGCTGATCGATAAGCTGAACTACATGGTGGACAAGAAGTCTAATCCTTGTGCAACAGGCGGCGCCCTGAAGGGCTATCAGATCACCAATAAGTTCGAGAGCTTTAAGTCCATGTCTACCCAGAACGGCTTCATCTTTTACATCCCTGCCTGGCTGACATCCAAGATCGATCCATCTACCGGCTTTGTGAACCTGCTGAAAACCAAGTATACCAGCATCGCCGATTCCAAGAAGTTCATCAGCTCCTTTGACAGGATCATGTACGTGCCCGAGGAGGATCTGTTCGAGTTTGCCCTGGACTATAAGAACTTCTCTCGCACAGACGCCGATTACATCAAGAAGTGGAAGCTGTACTCCTACGGCAACCGGATCAGAATCTTCCGGAATCCTAAGAAGAACAACGTGTTCGACTGGGAGGAGGTGTGCCTGACCAGCGCCTATAAGGAGCTGTTCAACAAGTACGGCATCAATTATCAGCAGGGCGATATCAGAGCCCTGCTGTGCGAGCAGTCCGACAAGGCCTTCTACTCTAGCTTTATGGCCCTGATGAGCCTGATGCTGCAGATGCGGAACAGCATCACAGGCCGCACCGACGTGGATTTTCTGATCAGCCCTGTGAAGAACTCCGACGGCATCTTCTACGATAGCCGGAACTATGAGGCCCAGGAGAATGCCATCCTGCCAAAGAACGCCGACGCCAATGGCGCCTATAACATCGCCAGAAAGGTGCTGTGGGCCATCGGCCAGTTCAAGAAGGCCGAGGACGAGAAGCTGGATAAGGTGAAGATCGCCATCTCTAACAAGGAGTGGCTGGAGTACGCCCAGACCAGCGTGAAGCACAAAAGGCCGGCGGCCACGAAAAAGGCCGGCCAGGCAAAAAAGAAAAAGGGATCCTACCCATACGATGTTCCAGATTACGCTTATCCCTACGACGTGCCTGATTATGCATACCCATATGATGTCCCCGACTATGCCTAAAGCCTCGAGGGTGGTGGTAGCGATTATAAAGATGATGATGATAAAGGCCTGAACGATATCTTTGAAGCCCAGAAAATTGAATGGCACGAGTAAAAGCTTGTCGAGCACCACCACCACCACCACTGAGATCCGGCTGCTAACAAAGCCCGAAAGGAAGCTGAGTTGGCTGCTGCCACCGCTGAGCAATAACTAGCATAACCCCTTGGGGCCTCTAAACGGGTCTTGAGGGGTTTTTTGCTGAAAGGAGGAACTATATCCGGATTGGCGAATGGGACGCGCCCTGTAGCGGCGCATTAAGCGCGGCGGGTGTGGTGGTTACGCGCAGCGTGACCGCTACACTTGCCAGCGCCCTAGCGCCCGCTCCTTTCGCTTTCTTCCCTTCCTTTCTCGCCACGTTCGCCGGCTTTCCCCGTCAAGCTCTAAATCGGGGGCTCCCTTTAGGGTTCCGATTTAGTGCTTTACGGCACCTCGACCCCAAAAAACTTGATTAGGGTGATGGTTCACGTAGTGGGCCATCGCCCTGATAGACGGTTTTTCGCCCTTTGACGTTGGAGTCCACGTTCTTTAATAGTGGACTCTTGTTCCAAACTGGAACAACACTCAACCCTATCTCGGTCTATTCTTTTGATTTATAAGGGATTTTGCCGATTTCGGCCTATTGGTTAAAAAATGAGCTGATTTAACAAAAATTTAACGCGAATTTTAACAAAATATTAACGCTTACAATTTAGGTGGCACTTTTCGGGGAAATGTGCGCGGAACCCCTATTTGTTTATTTTTCTAAATACATTCAAATATGTATCCGCTCATGAGACAATAACCCTGATAAATGCTTCAATAATATTGAAAAAGGAAGAGTATGAGTATTCAACATTTCCGTGTCGCCCTTATTCCCTTTTTTGCGGCATTTTGCCTTCCTGTTTTTGCTCACCCAGAAACGCTGGTGAAAGTAAAAGATGCTGAAGATCAGTTGGGTGCACGAGTGGGTTACATCGAACTGGATCTCAACAGCGGTAAGATCCTTGAGAGTTTTCGCCCCGAAGAACGTTTTCCAATGATGAGCACTTTTAAAGTTCTGCTATGTGGCGCGGTATTATCCCGTATTGACGCCGGGCAAGAGCAACTCGGTCGCCGCATACACTATTCTCAGAATGACTTGGTTGAGTACTCACCAGTCACAGAAAAGCATCTTACGGATGGCATGACAGTAAGAGAATTATGCAGTGCTGCCATAACCATGAGTGATAACACTGCGGCCAACTTACTTCTGACAACGATCGGAGGACCGAAGGAGCTAACCGCTTTTTTGCACAACATGGGGGATCATGTAACTCGCCTTGATCGTTGGGAACCGGAGCTGAATGAAGCCATACCAAACGACGAGCGTGACACCACGATGCCTGCAGCAATGGCAACAACGTTGCGCAAACTATTAACTGGCGAACTACTTACTCTAGCTTCCCGGCAACAATTAATAGACTGGATGGAGGCGGATAAAGTTGCAGGACCACTTCTGCGCTCGGCCCTTCCGGCTGGCTGGTTTATTGCTGATAAATCTGGAGCCGGTGAGCGTGGTAGTCGCGGTATCATTGCAGCACTGGGGCCAGATGGTAAGCCCTCCCGTATCGTAGTTATCTACACGACGGGGAGTCAGGCAACTATGGATGAACGAAATAGACAGATCGCTGAGATAGGTGCCTCACTGATTAAGCATTGGTAACTGTCAGACCAAGTTTACTCATATATACTTTAGATTGATTTAAAACTTCATTTTTAATTTAAAAGGATCTAGGTGAAGATCCTTTTTGATAATCTCATGACCAAAATCCCTTAACGTGAGTTTTCGTTCCACTGAGCGTCAGACCCCGTAGAAAAGATCAAAGGATCTTCTTGAGATCCTTTTTTTCTGCGCGTAATCTGCTGCTTGCAAACAAAAAAACCACCGCTACCAGCGGTGGTTTGTTTGCCGGATCAAGAGCTACCAACTCTTTTTCCGAAGGTAACTGGCTTCAGCAGAGCGCAGATACCAAATACTGTCCTTCTAGTGTAGCCGTAGTTAGGCCACCACTTCAAGAACTCTGTAGCACCGCCTACATACCTCGCTCTGCTAATCCTGTTACCAGTGGCTGCTGCCAGTGGCGATAAGTCGTGTCTTACCGGGTTGGACTCAAGACGATAGTTACCGGATAAGGCGCAGCGGTCGGGCTGAACGGGGGGTTCGTGCACACAGCCCAGCTTGGAGCGAACGACCTACACCGAACTGAGATACCTACAGCGTGAGCTATGAGAAAGCGCCACGCTTCCCGAAGGGAGAAAGGCGGACAGGTATCCGGTAAGCGGCAGGGTCGGAACAGGAGAGCGCACGAGGGAGCTTCCAGGGGGAAACGCCTGGTATCTTTATAGTCCTGTCGGGTTTCGCCACCTCTGACTTGAGCGTCGATTTTTGTGATGCTCGTCAGGGGGGCGGAGCCTATGGAAAAACGCCAGCAACGCGGCCTTTTTACGGTTCCTGGCCTTTTGCTGGCCTTTTGCTCACATGTTCTTTCCTGCGTTATCCCCTGATTCTGTGGATAACCGTATTACCGCCTTTGAGTGAGCTGATACCGCTCGCCGCAGCCGAACGACCGAGCGCAGCGAGTCAGTGAGCGAGGAAGCGGAAGAGCGCCTGATGCGGTATTTTCTCCTTACGCATCTGTGCGGTATTTCACACCGCAATGGTGCACTCTCAGTACAATCTGCTCTGATGCCGCATAGTTAAGCCAGTATACACTCCGCTATCGCTACGTGACTGGGTCATGGCTGCGCCCCGACACCCGCCAACACCCGCTGACGCGCCCTGACGGGCTTGTCTGCTCCCGGCATCCGCTTACAGACAAGCTGTGACCGTCTCCGGGAGCTGCATGTGTCAGAGGTTTTCACCGTCATCACCGAAACGCGCGAGGCAGCTGCGGTAAAGCTCATCAGCGTGGTCGTGAAGCGATTCACAGATGTCTGCCTGTTCATCCGCGTCCAGCTCGTTGAGTTTCTCCAGAAGCGTTAATGTCTGGCTTCTGATAAAGCGGGCCATGTTAAGGGCGGTTTTTTCCTGTTTGGTCACTGATGCCTCCGTGTAAGGGGGATTTCTGTTCATGGGGGTAATGATACCGATGAAACGAGAGAGGATGCTCACGATACGGGTTACTGATGATGAACATGCCCGGTTACTGGAACGTTGTGAGGGTAAACAACTGGCGGTATGGATGCGGCGGGACCAGAGAAAAATCACTCAGGGTCAATGCCAGCGCTTCGTTAATACAGATGTAGGTGTTCCACAGGGTAGCCAGCAGCATCCTGCGATGCAGATCCGGAACATAATGGTGCAGGGCGCTGACTTCCGCGTTTCCAGACTTTACGAAACACGGAAACCGAAGACCATTCATGTTGTTGCTCAGGTCGCAGACGTTTTGCAGCAGCAGTCGCTTCACGTTCGCTCGCGTATCGGTGATTCATTCTGCTAACCAGTAAGGCAACCCCGCCAGCCTAGCCGGGTCCTCAACGACAGGAGCACGATCATGCGCACCCGTGGGGCCGCCATGCCGGCGATAATGGCCTGCTTCTCGCCGAAACGTTTGGTGGCGGGACCAGTGACGAAGGCTTGAGCGAGGGCGTGCAAGATTCCGAATACCGCAAGCGACAGGCCGATCATCGTCGCGCTCCAGCGAAAGCGGTCCTCGCCGAAAATGACCCAGAGCGCTGCCGGCACCTGTCCTACGAGTTGCATGATAAAGAAGACAGTCATAAGTGCGGCGACGATAGTCATGCCCCGCGCCCACCGGAAGGAGCTGACTGGGTTGAAGGCTCTCAAGGGCATCGGTCGAGATCCCGGTGCCTAATGAGTGAGCTAACTTACATTAATTGCGTTGCGCTCACTGCCCGCTTTCCAGTCGGGAAACCTGTCGTGCCAGCTGCATTAATGAATCGGCCAACGCGCGGGGAGAGGCGGTTTGCGTATTGGGCGCCAGGGTGGTTTTT
